# Endothelial metastasis-associated protein 1 (MTA1) is an essential molecule for angiogenesis

**DOI:** 10.1101/2021.04.15.439980

**Authors:** Mizuho Ishikawa, Mitsuhiko Osaki, Narumi Uno, Takahito Ohira, Hiroyuki Kugoh, Futoshi Okada

**Author notes:** **Address correspondence to:** Mitsuhiko Osaki, Ph.D. Division of Experimental Pathology, Faculty of Medicine, Tottori University, 86, Nishi-cho, Yonago, Tottori 683-8503, Japan.

## Abstract

Our previous study revealed that metastasis-associated protein 1 (MTA1), expressed in endothelial cells acts as a factor promoting tube formation. In this study, we aimed to clarify the importance of MTA1 in tube formation using *MTA1*-knockout endothelial cells (MTA1-KO MSS31 cells). Tube formation was strongly suppressed in MTA1-KO MSS31 cells, whereas MTA1-overexpressed MTA1-KO MSS31 cells regained the ability to form tube-like structures. Furthermore, MTA1-KO MSS31 cells showed higher phosphorylation of non-muscle myosin heavy chain IIa (NMIIa), which resulted in suppression of tube formation; this was attributed to a decrease of MTA1– S100 calcium-binding protein A4 (S100A4) complex formation. Moreover, inhibition of tube formation in MTA1-KO MSS31 cells could not be rescued by stimulation with vascular endothelial growth factor (VEGF). These results demonstrated that MTA1 is an essential molecule for angiogenesis, and that it may be involved in different steps of the angiogenic process compared to the VEGF–VEGF receptor 2 (VEGFR2) pathway. Our findings showed that endothelial MTA1 and its pathway are promising targets for inhibiting tumor angiogenesis, supporting the development of MTA1-based anti-angiogenic therapies.

## Introduction

Metastasis-associated protein 1 (MTA1), a member of the MTA family, was first identified by Toh and Nicolson (Toh et al, 1994). MTA1 expression is reported to be correlated with tumor malignancy in several cancer types, including esophageal cancer, gastrointestinal cancer, non–small-cell lung cancer, breast cancer, and ovarian cancer (Toh et al, 1997; Toh et al, 1999; Zhu et al, 2010; Jang et al, 2006; Dannenmann et al, 2008). Furthermore, MTA1 in tumor cells promotes proliferation, invasion, tumor angiogenesis, and metastasis (Nicolson et al, 2003; Kai et al, 2011; Pakala et al, 2013).

However, the precise functional role of endothelial MTA1 in angiogenesis remains unclear. In a previous study, we examined the function of MTA1 in endothelial cells by *MTA1* knockdown and found that MTA1 small interfering RNA (siRNA) significantly inhibits tube formation but not the proliferation and migration of endothelial cells. Moreover, we demonstrated that *MTA1* knockdown induced a decrease in S100 calcium-binding protein A4 (S100A4) expression and an increase in phosphorylated non-muscle myosin heavy chain IIa (NMIIa). This phosphorylation level of NMIIa may influence the formation of tube structures via altered cytoskeletal dynamics in endothelial cells (Ishikawa et al, 2019).

In the present study, *MTA1*-knockout endothelial cells (MTA1-KO MSS31 cells) were established and examined to clarify the role of MTA1 expression in tube formation by endothelial cells. We also investigated the relationship between the inhibition of tube formation in MTA1-KO MSS31 cells and the vascular endothelial growth factor (VEGF) –vascular endothelial growth factor receptor 2 (VEGFR2) pathway.

## Results

### Engineering of MTA1-KO MSS31 clones

To gain deeper insight into the functions of *MTA1*, we designed gRNA targeting *MTA1* and used the CRISPR-Cas9 system to generate MTA1-KO MSS31 cell lines. We generated two colonies (#1 and #2) grown from individual cells sorted by fluorescence-activated cell sorting (FACS). These colonies were confirmed as *MTA1* knockout by western blotting (Fig EV1A). We also checked the deletion of *MTA1* in the cellular genome of the two colonies by genome sequencing. Clone #1 had a four-base deletion in one allele and a 116-base deletion. Clone #2 had a five-base deletion in one allele and a 201-base deletion (Fig EV1B). Frameshift mutation due to multiple-base deletion in each allele resulted in the loss of wild-type MTA1 protein expression. These results indicated the successful generation of MTA1-KO MSS31 cell clones.

### MTA1 expression contributes to tube formation in vitro

Previously, we reported that suppression of MTA1 in endothelial cells inhibited tube formation *in vitro* (Ishikawa et al, 2019). We then performed a tube formation assay using wild-type MSS31 cells and two MTA1-KO clones to determine the effect of tube formation in MTA1-KO clones. When we observed tube formation at 5 h after the cells were plated onto Geltrex, wild-type MSS31 cells showed tube formation spanning the entire well. In contrast, two MTA1-KO clones showed significantly impaired tube formation (Fig 1A, B).

**Fig 1.**
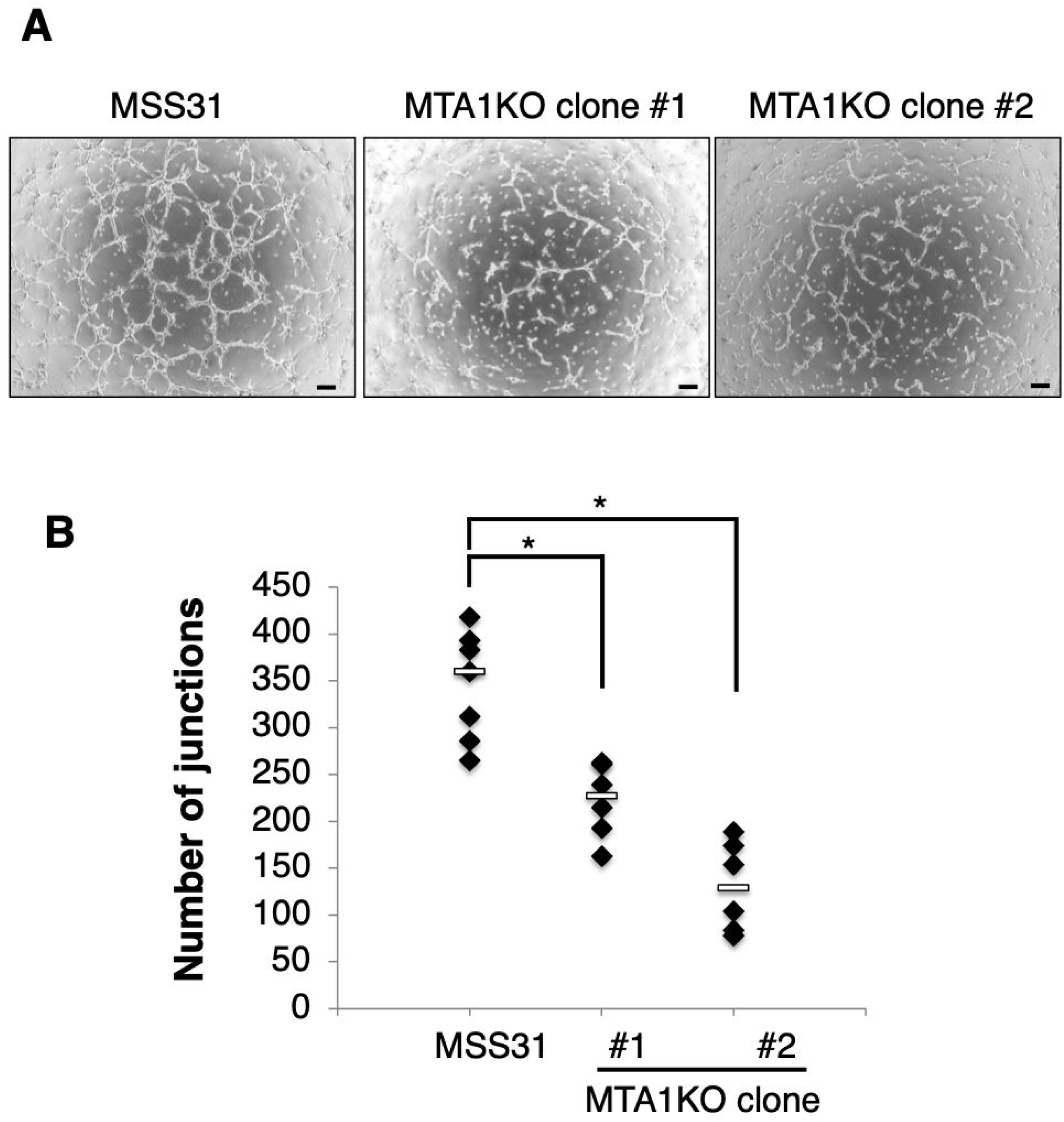
MTA1-KO clones showed inhibition of tube formation at 5 h after seeding wild-type MSS31 cells and two MTA1-KO clones. (A)Wild-type MSS31 cells and two MTA1-KO clones were seeded on Geltrex, and tube formation was photographed after 5 h. Scale bars, 200 μm. (B) Quantification of the number of junctions in wild-type MSS31 cells and MTA1-KO clones (wild-type MSS31 cells: n = 7, MTA1-KO clones: n = 6). *p < 0.05 compared to the wild-type MSS31 cells

We also hypothesized that diminished tube formation in MTA1-KO clones could be recovered by MTA1 overexpression. After MTA1 overexpression in MTA1-KO MSS31 clones transfected with MTA1 expression vector (MTA1-KO+OE MSS31 clones: OE) was confirmed by real-time PCR and western blotting (Fig EV2A, B), we performed a tube formation assay using MTA1-KO clones and MTA1-KO+OE MSS31 clones. At 19 h after the cells were plated onto Geltrex, short and incomplete tube-like structures were observed in MTA1-KO+OE MSS31 clones; however, no tube formation was observed in MTA1-KO clones that were transfected with empty vector or remained untransfected (Fig 2A, B). These results indicated that the inhibition of tube formation caused by *MTA1* knockout was recovered by MTA1 overexpression.

**Fig 2.**
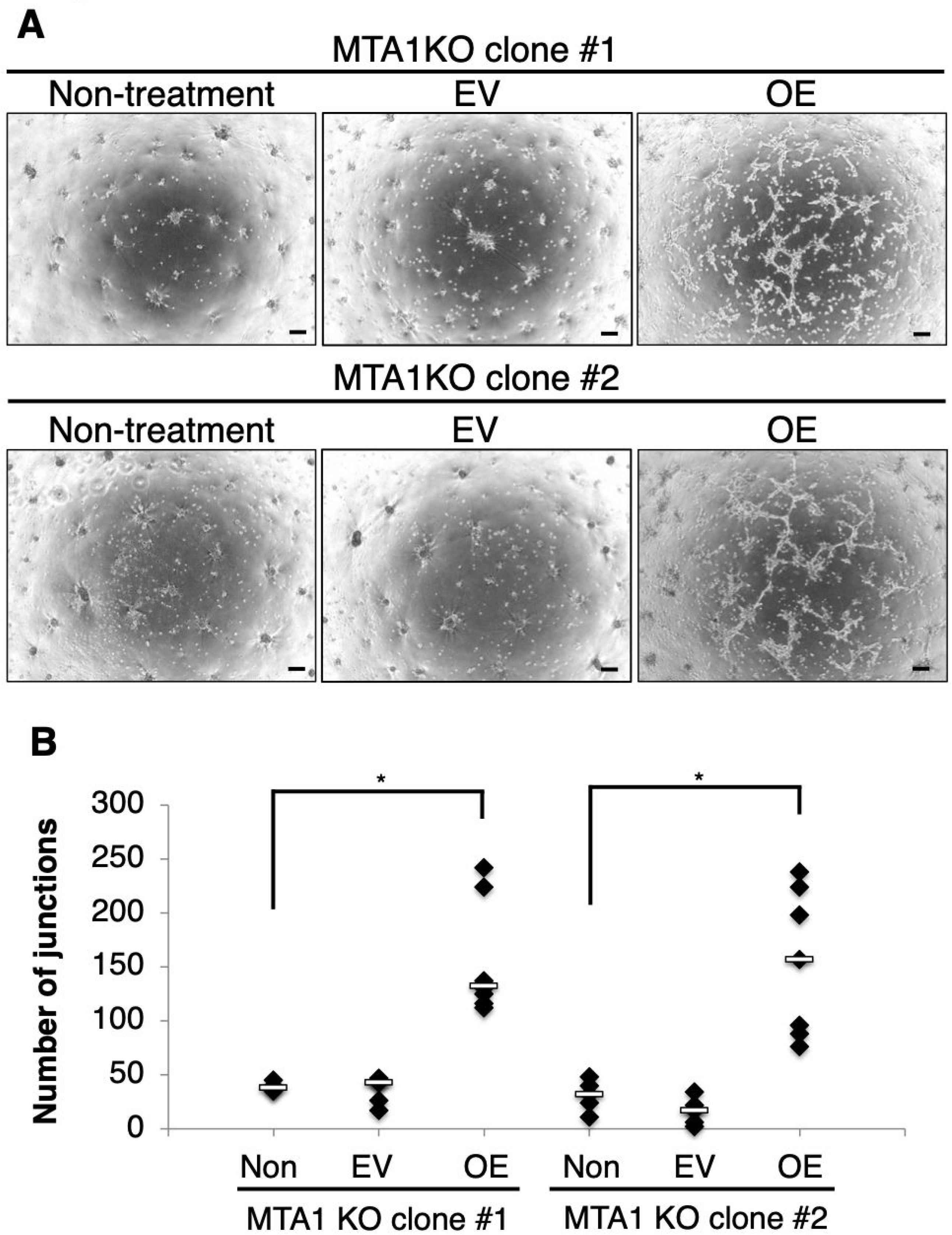
MTA1-KO MSS31 cells transfected with MTA1 expression vector (MTA1-KO+OE MSS31 cells) can partly recover tube formation. (A) Tube formation assays were performed using MTA1-KO MSS31 cells and MTA1-KO+OE MSS31 clones and were photographed after 19 h. Scale bars, 200 μm. (B) Quantification of the number of junctions in MTA1-KO MSS31 cells (Non-treatment), MTA1-KO cells transfected with empty vector (EV), and MTA1-KO cells transfected with MTA1 expression vector (MTA1-KO+OE MSS31 cells: OE) (Non-treatment: n = 6, EV: n = 6, OE: n = 8). *p < 0.05 compared to the non-treated counterpart of each clone

### Involvement of MTA1–S100A4 interaction and phosphorylated NMIIa in tube formation in MTA1-KO and MTA1-KO+OE MSS31 cells

We previously reported the mechanism of tube formation by MTA1, wherein MTA1 interacts with S100A4 and changes the phosphorylation state of NMIIa (Ishikawa et al, 2019). Therefore, we examined whether inhibition of tube formation by *MTA1* knockout and its restoration by MTA1 overexpression was correlated with the amount of MTA1–S100A4 complex and the phosphorylation state of NMIIa. First, MTA1–S100A4 interaction was found to be disappeared in MTA1-KO MSS31 cells and was increased in MTA1-KO+OE MSS31 cells compared to the extent of MTA1– S100A4 interaction in MTA1-KO MSS31 cells; this was evaluated by immunoprecipitation–western blot using anti-MTA1 or anti-S100A4 antibodies (Fig 3A, B). Second, the phosphorylation of NMIIa was found to be increased by *MTA1* knockout (Fig 4A, B). The increased NMIIa phosphorylation was slightly impeded by MTA1 overexpression; however, the difference was not statistically significant (Fig 4A, B). Thus, we proposed that the altered tube formation abilities of *MTA1*-knockout and -overexpressing cells were caused by MTA1–S100A4 interaction and NMIIa phosphorylation.

**Fig 3.**
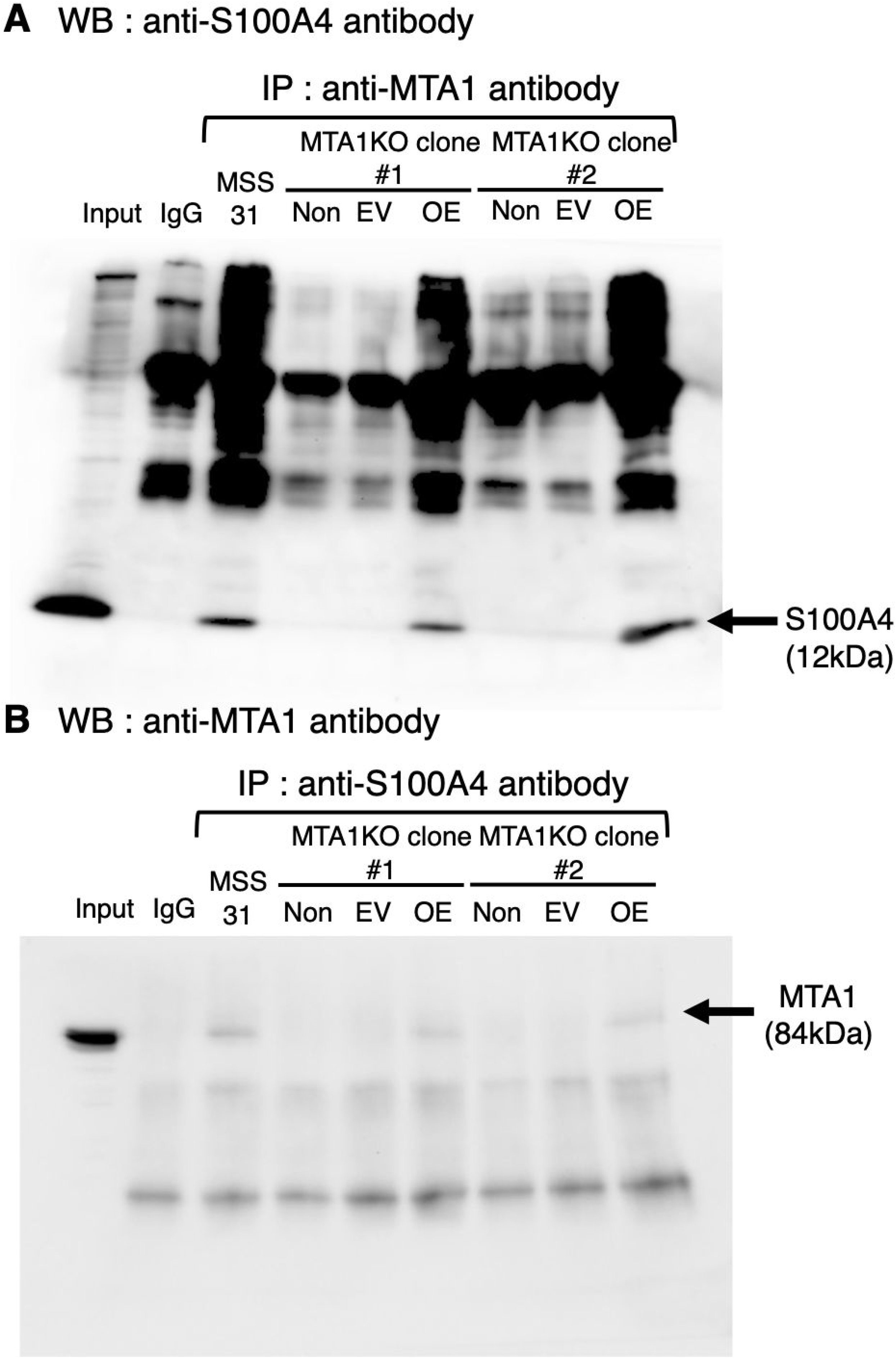
Formation of the MTA1–S100A4 complex in MTA1-KO MSS31 cells and in MTA1-KO MSS31 cells transfected with the MTA1 expression vector (MTA1-KO+OE MSS31 cells). (A) Western blotting with anti-S100A4 antibody after IP with the indicated antibodies using wild-type MSS31 cells, MTA1-KO MSS31 cells (Non-treatment), MTA1-KO cells transfected with empty vector (EV), and MTA1-KO cells transfected with MTA1 expression vector (MTA1-KO+OE MSS31 cells: OE). Input represents 10% of the cell lysate. Normal rabbit IgG was used as a negative control (n = 3). (B) Western blotting with anti-MTA1 antibody after IP with the indicated antibodies using cell lysates. Normal mouse IgG was used as a negative control (n = 4) IP: immunoprecipitation

**Fig 4.**
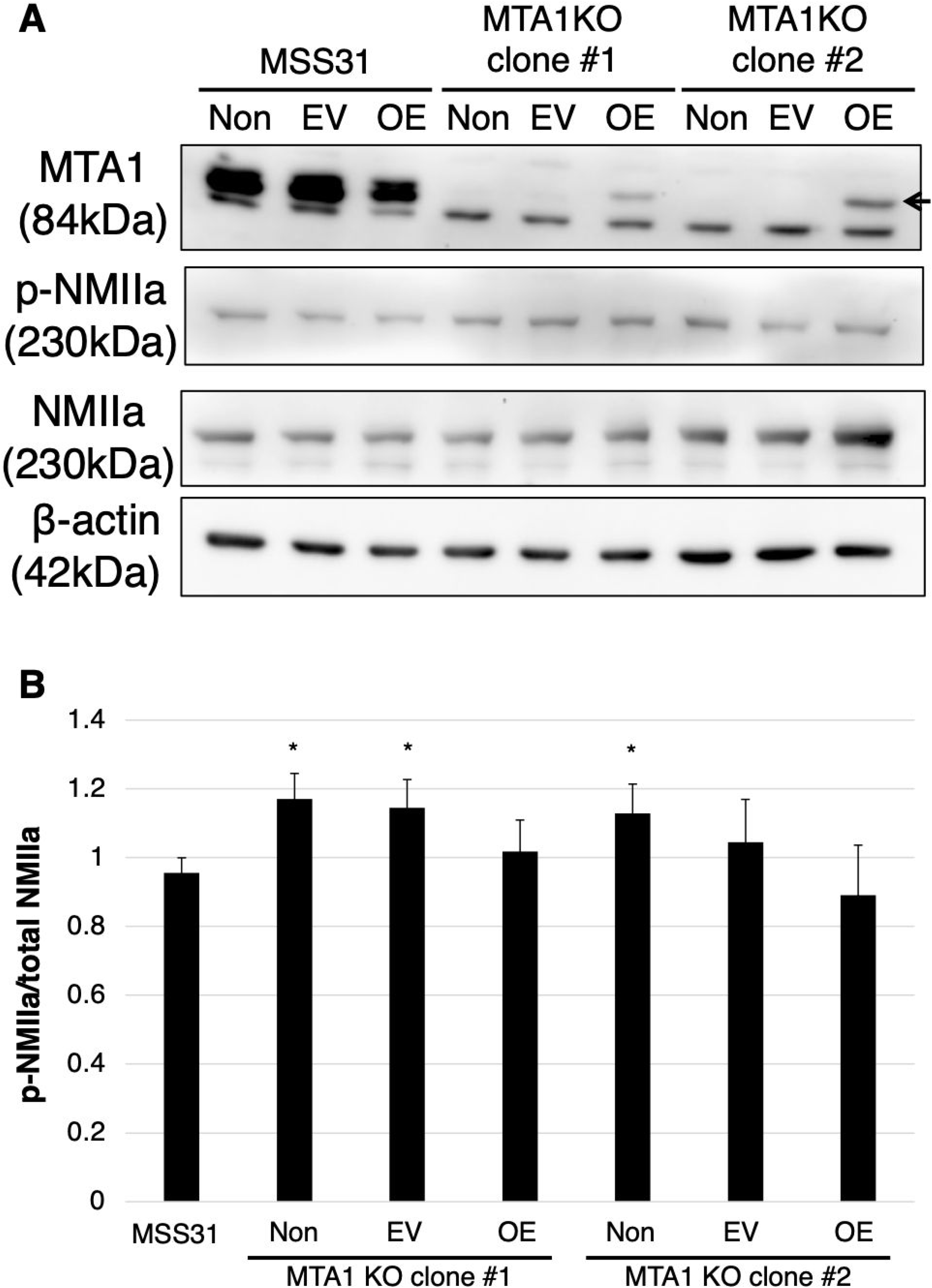
Changes in the phosphorylation status of NMIIa in wild-type MSS31 cells and in MTA1-KO MSS31 cells transfected with the MTA1 expression vector. (A) Western blotting of wild-type MSS31 cells, MTA1-KO MSS31 cells (Non-treatment), MTA1-KO cells transfected with empty vector (EV), and MTA1-KO cells transfected with MTA1 expression vector (MTA1-KO+OE MSS31 cells: OE) (n = 6). (B) Phosphorylated NMIIa (p-NMIIa) expression levels were measured using ImageJ software and normalized to the expression of β-actin (n = 6). *p < 0.05 compared to the wild-type MSS31 cells

### Relationship between MTA1 and VEGF in MTA1-KO MSS31 cells

We revealed that *MTA1* knockdown and knockout inhibited tube formation *in vitro*. Angiogenesis is influenced by numerous angiogenic pathways, including the VEGF–VEGFR2 pathway. Therefore, we examined whether the suppressed tube formation in MTA1-KO MSS31 cells could be rescued by stimulation with VEGF. We first confirmed that VEGFR2 was phosphorylated by VEGF stimulation in both wild-type MSS31 cells and MTA1-KO MSS31 cells; furthermore, VEGFR2 expression levels were not altered by *MTA1* knockout (Fig EV3). Tube formation was promoted in wild-type MSS31 cells treated with VEGF. In contrast, despite the phosphorylation of VEGFR2 by VEGF treatment, tube formation could not be rescued in MTA1-KO MSS31 cells treated with VEGF (Fig 5A, B and Fig EV3). Furthermore, MTA1 expression was not affected by VEGF stimulation (Fig EV4). These results indicated that VEGF stimulation could not promote tube formation in MTA1-KO MSS31 cells, implying that the role of MTA1 in tube formation may be independent of VEGF.

**Fig 5.**
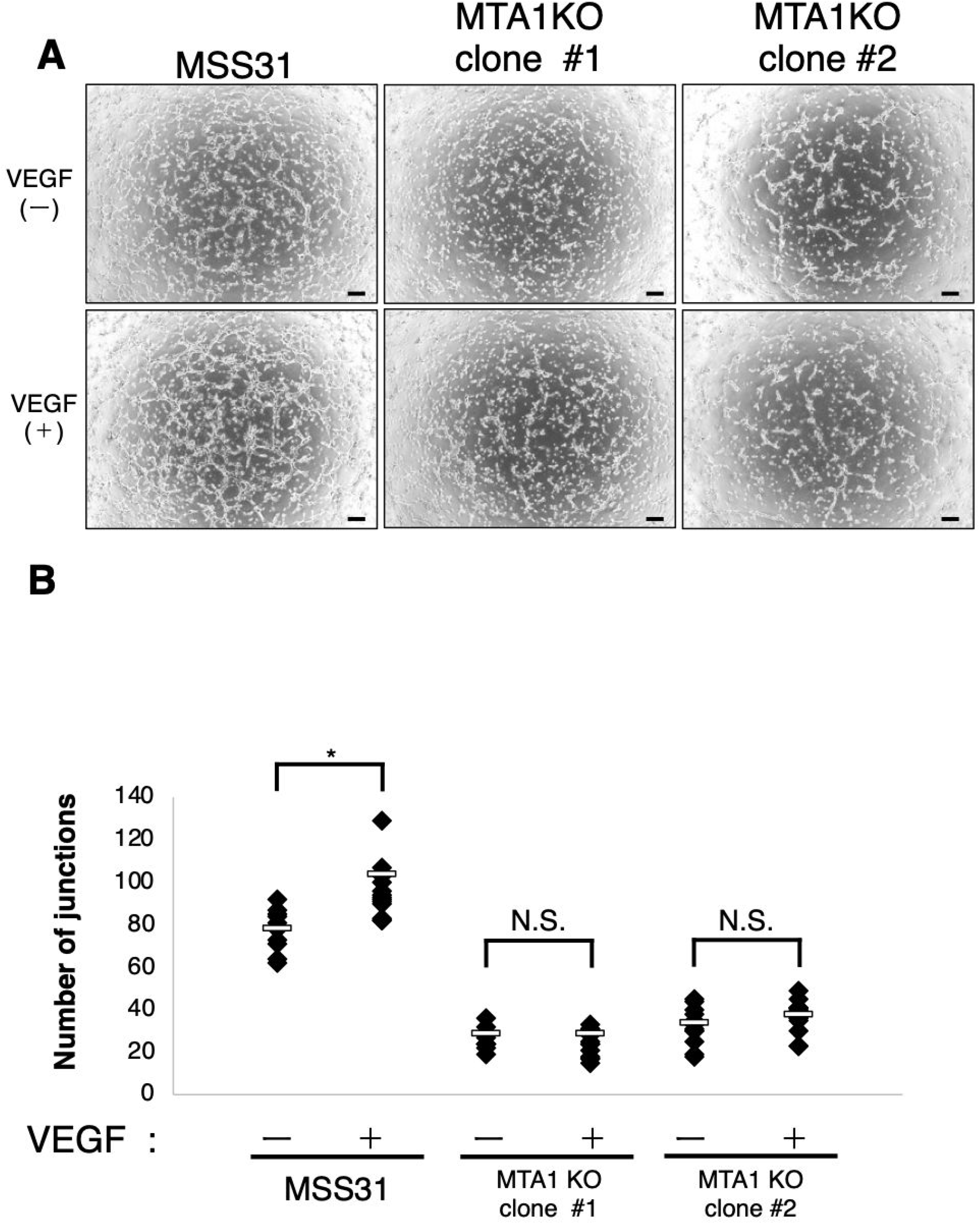
The interaction of MTA1 and VEGF in MTA1-KO MSS31 cells. (A) Tube formation assays were performed using wild-type MSS31 cells and MTA1-KO clones, which were treated with VEGF/left untreated, and were photographed after 4 h. Scale bars, 200 μm. (B) Quantification of the number of junctions of wild-type MSS31 cells and MTA1-KO clones, which were treated with VEGF or not (VEGF (-): n = 16, VEGF (+): n = 20). *p < 0.05 compared to wild-type MSS31 cells without VEGF treatment or each MTA1-KO MSS31 cell clone without VEGF treatment

## Discussion

MTA1 expression in tumor cells is reported to exert multiple functions involved in cancer progression such as proliferation, migration, invasion, and tumor angiogenesis (Nicolson et al, 2003; Kai et al, 2011; Pakala et al, 2013). However, the role of MTA1 in endothelial cells is not clearly elucidated. Our group previously reported that MTA1 is involved in tumor angiogenesis via the MTA1–S100A4–NMIIa axis in endothelial cells (Ishikawa et al, 2019). Therefore, we attempted to confirm the function of endothelial MTA1 in angiogenesis using MTA1-KO MSS31 cells generated with the CRISPR-Cas9 system.

The phosphorylation level of NMIIa decreased when MTA1 was overexpressed in the MTA1-KO MSS31 cells compared to the corresponding phosphorylation level in MTA1-KO MSS31 cells and MTA1-KO cells transfected with empty vector (Fig 4B) Although the protein-expression level of MTA1 in MTA1-KO+OE MSS31 cells was lower than that in the wild-type MSS31 cells (Fig 4A), the interaction between MTA1 and S100A4 occurs similar to that seen in the wild-type MSS31 cells (Fig 3A, B). These results indicated that even a minute concentration of MTA1 was sufficient to impact angiogenesis via the interaction with S100A4.

We observed that MTA1-KO MSS31 cells showed impaired tube formation at 5 h after the cells were seeded on Geltrex (Fig 1A, B) and the tube structure disappeared at 19 h (Fig 2A, B). In contrast, MTA1-KO MSS31 cells transfected with the MTA1 expression vector (MTA1-KO+OE MSS31 cells) partially formed tube-like structures at 19 h after the cells were seeded on Geltrex (Fig 2A, B). Tube formation in MTA1-KO+OE MSS31 cells was partially observed at 19 h, but not at 5 h, possibly because the direct use of MTA1 expression vector-transfected cells in the tube formation assay caused differences in MTA1 expression levels among the MSS31 cells. These results indicated that endothelial MTA1 may play an important role in tube formation as well as in maintenance.

Angiogenesis involves multiple steps including protease production, endothelial migration and proliferation, vascular tube formation, and blood vessel maturation, which are regulated by numerous angiogenic pathways (Rajabi & Mousa, 2017; Herbert & Stainier, 2011; Welti et al, 2013). This process can be roughly divided into two steps: degradation of the basement membrane, proliferation, migration, and sprout formation (Step 1); and vascular tube formation and maturation (Step 2) (Fig 6). Since VEGF– VEGFR2 signaling is a key pathway in physiological and tumor angiogenesis (Ferrara et al, 2003), we examined the correlation between MTA1 and VEGF *in vitro*. As shown in Figure 5, the tube formation ability was not recovered regardless of whether MTA1-KO MSS31 cells were treated with VEGF or not, although the phosphorylation of VEGFR2 was confirmed in those cells treated with VEGF (Fig EV3). These data indicated that the VEGF–VEGFR2 signaling pathway controls the survival, migration, and proliferation of vascular endothelial cells in the first step of the angiogenic process (Ferrara et al, 2003). Conversely, the MTA1–S100A4–NMIIa axis primarily regulates tube formation in the second step. In other words, it may be possible to inhibit angiogenesis by blocking “Step 2” with or without activation of “Step 1”.

**Fig 6.**
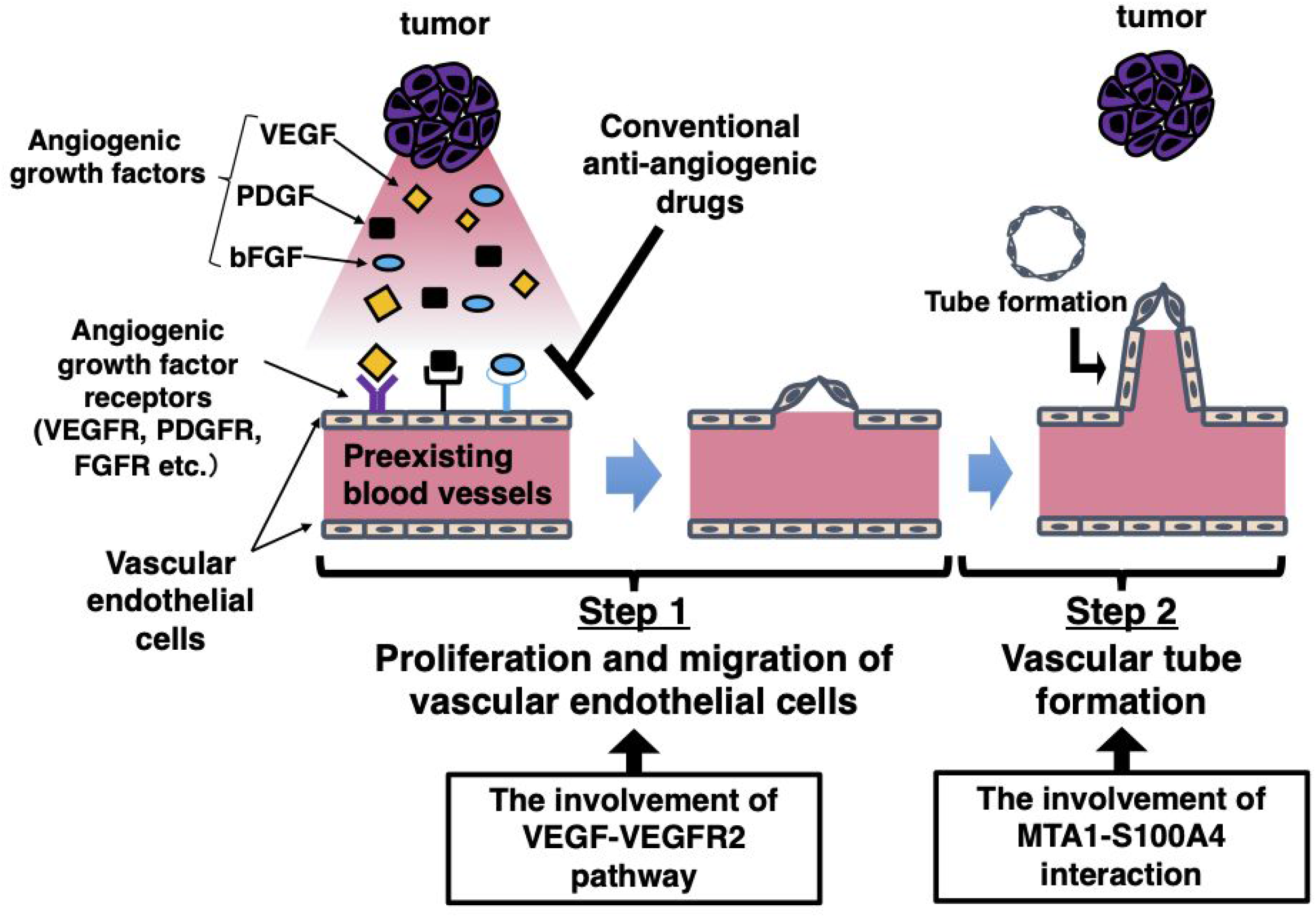
Schematic diagram of the process of tumor angiogenesis New blood vessels are generated from pre-existing blood vessels through angiogenic growth factors secreted by tumor cells. The vascular endothelial cells in pre-existing blood vessels degrade the basement membrane, proliferate, migrate, and form sprouts after the activation of angiogenic stimuli (Step 1). Subsequently, these sprouts lead to vascular tube formation, followed by blood vessel maturation to complete the tube structure through which blood can flow (Step 2). VEGF stimulates endothelial proliferation and migration in Step 1, whereas MTA1 is primarily correlated with vascular tube formation in Step 2

Tumor angiogenesis is an appealing therapeutic target (Folkman, 1971). Anti-angiogenic therapy has been applied clinically and has emerged as one of the viable treatment options for cancer (Jayson et al, 2016). Due to the pivotal role of the VEGF–VEGFR2 pathway in pathological angiogenesis, most approved anti-angiogenic drugs target VEGF-A or its receptors, including VEGFR2 (Ferrara & Kerbel, 2005; Haibe et al, 2020). Although these treatment strategies have provided substantial clinical benefit in cancer therapeutics, their effects are limited by the development of resistance (Bergers & Hanahan, 2008; Beijnum et al, 2015). In the present study, we revealed that the MTA1–S100A4 axis in endothelial cells is an important pathway for tube formation and is functionally distinct from the VEGF–VEGFR2 pathway at the site of action in angiogenesis. Thus, therapies targeting the endothelial MTA1–S100A4 axis may provide a novel option distinct from conventional therapies. In other words, the axis may serve as a useful target molecule for treating patients who manifest VEGF-resistance or show poor responses to VEGF inhibitors. Moreover, the combination of anti-angiogenic drugs targeting the VEGF–VEGFR2 pathway, as well as the MTA1–S100A4 pathway, may induce additive or synergistic effects in the inhibition of tumor angiogenesis and tumor growth, resulting in improved therapeutic outcomes.

Taken together, our data confirmed that MTA1 expression in endothelial cells plays a crucial role in tube formation; additionally, MTA1 is involved in a step that is distinct of the role of VEGF in the process of angiogenesis. Based on these findings, and the results of our previous study, the MTA1 or MTA1–S100A4–NMIIa pathway in endothelial cells could prove to be an attractive target for suppressing angiogenesis and tumor growth, with a mechanism of action different from that of VEGF inhibitors.

## Materials and Methods

### Cell culture and reagents

Mouse endothelial cells (MSS31 cells) and MTA1-KO MSS31 cells were maintained in Dulbecco’s modified Eagle’s medium (DMEM)/Ham’s F-12 medium supplemented with 10% fetal bovine serum (FBS). These cell lines were cultured at 37°C in a humidified 5% CO_2_ /95% air mixture. MSS31 cells and MTA1-KO MSS31 cells were transfected with the MTA1 expression vector gifted by Dr. K. Takenaga (Chiba Cancer Center Research Institute, Chiba, Japan) using Lipofectamine 2000 (Invitrogen, Carlsbad, CA, USA), in accordance with the manufacturer’s instructions.

### Engineering of guide RNA (gRNA) and CRISPR-Cas9 vector for MTA1 knockout

The guide RNA (gRNA) for *MTA1* knockout targeting the first exon of *MTA1* on the coding strand was engineered using the online CRISPR design tool (http://crispr.mit.edu/). The targeted sequence was designed 5’-CAGGATTGAAGAGCTTAACAAGG-3’. Complementary guide oligonucleotides (forward: 5’-CACCGCAGGATTGAAGACCTTAACA-3’; reverse: 5’-AAACTGTTAAGCTCTTCAATCCTGC-3’) were custom synthesized separately by Sigma-Aldrich (St. Louis, MO, USA), annealed, and cloned into the BbsI site of pSpCas9(BB)-2A-GFP (pX458; Addgene, Cambridge, MA, USA). PX 458 was a gift from Feng Zhang (Addgene plasmid # 48138 ; http://n2t.net/addgene:48138 ; RRID: Addgene_48138). (Ref: PubMed 24157548)

### Electroporation and cell sorting

MSS31 cells were harvested and counted. Thereafter, 1−2 × 10^6^ cells were resuspended with 3 μg plasmid DNA in 100 mL of electroporation buffer, transferred to a 0.2 cm cuvette (Nepa Gene, Japan), and electroporated using a square electric pulse generating electroporator, NEPA21 (Nepa Gene, Japan), i.e., Poring Pulse (voltage: 150 V, pulse length: 7.5 msec, pulse interval: 50 msec, number of pulses: 2, Decay Rate 10%, polarity: +), Transfer Pulse (voltage: 20 V, pulse length: 50 msec, pulse interval: 50 msec, number of pulses: 5, polarity: +/-). The cells were transferred to complete medium and seeded in 60 mm dishes. After 24 h of electroporation, the transfected populations were sorted by GFP expression using fluorescence-activated cell sorting (FACS). Single cells were sorted into 384-well microplates and cultured for more than a month to permit colony formation.

### DNA sequencing

PCR products were directly sequenced using specific primers and were cloned into the pCR™4-TOPO® TA vector (Thermo Fisher Scientific, Rochester, NY, USA) using the TOPO™ TA Cloning™ Kit for Sequencing, without competent cells (Thermo Fisher Scientific), following the manufacturer’s instructions. Plasmid DNA was isolated using the PureYield™ Plasmid Miniprep System (Promega, Madison, WI, USA).

Plasmids were sequenced using the M13 forward primer (5’-GTAAAACGACGGCCAG-3’) and M13 reverse primer (5’-CAGGAAACAGCTATGAC-3’).

### Tube formation assay

For the tube formation assays, 24-well plates were coated with Geltrex, which was allowed to solidify overnight at 4 °C. MSS31 cells and MTA1-KO MSS31 cells (8 × 10^4^ cells/well) were seeded and cultured in serum-free medium or serum-free medium supplemented with VEGF-A (10 ng/mL). Tube formation was evaluated after 4 h, 5 h, or 19 h; subsequently, representative fields (4× magnification) were photographed with a KEYENCE BZ-X710 microscope (Keyence, Osaka, Japan). The number of junctions, branches, and total branching length were quantified using an angiogenesis analyzer (developed by Gilles Carpentier) in Image J (National Institute of Health, Bethesda, MD, USA).

### Western blotting

Cells were washed in cold PBS and lysed in lysis buffer containing 20 mM Tris-HCl (pH 7.4), 150 mM NaCl, 0.1% sodium dodecyl sulfate (SDS), 1% sodium deoxycholate, 1% Triton X-100, 1 μg/mL aprotinin, and 1 μg/mL leupeptin. Lysates were centrifuged at 15,000 rpm for 5 min. Protein concentrations were estimated using the Bradford protein assay (Bio-Rad Laboratories, Hercules, CA, USA) with bovine serum albumin (BSA) as the standard. Proteins were resolved by SDS-polyacrylamide gel electrophoresis using 8%, 10%, 12%, and 15% gels, following which they were electrotransferred to polyvinylidene difluoride (PVDF) membranes (Millipore, Bedford, MA, USA). The membranes were then blotted using primary antibodies, washed, and incubated with secondary antibodies. The membranes were washed, following which the bound antibodies were detected using an enhanced chemiluminescence detection system (Amersham, Buckinghamshire, UK). The primary antibodies used in the present study are listed as follows: anti-MTA1 polyclonal antibody (1:2000 dilution; Abcam, Cambridge, UK), anti-S100A4 polyclonal antibody (1:1000 dilution; Millipore), anti-NMIIA polyclonal antibody (1:1000 dilution; Cell Signaling Technology, Danvers, MA, USA), and anti-phospho-NMIIA polyclonal antibody (1:1000 dilution; Cell Signaling Technology). The secondary antibodies used in the present study are listed as follows: anti-mouse IgG-HRP (MBL, Nagoya, Japan) and goat anti-rabbit IgG-horseradish peroxidase (HRP; Santa Cruz Biotechnology, Santa Cruz, CA, USA). Each experiment using western blotting was replicated three or four times.

### Real-time PCR analysis

Total RNA was extracted using TRIzol reagent (Thermo Fisher Scientific, Rochester, NY, USA), and 1 μg of total RNA was used to produce cDNA using the TaKaRa PrimeScript RT master mix (TaKaRa Bio, Otsu, Japan). Two microliters of cDNA was then used for quantitative PCR. Quantitative real-time PCR was performed on a Roche LightCycler 480 (Roche Diagnostics, Palo Alto, CA, USA) using SYBR Premix Ex Taq II (Tli RNaseH Plus; TaKaRa Bio). The primer sequences are as follows: *MTA1* qRT forward primer (5’-GCGGCGAATGAACTGGA-3’) and qRT reverse primer (5’-TTGGTTTCTGAGGATGAGAGCA-3’). The reaction mixtures were incubated at 95°C for 30 s, then at 95°C for 10 s, followed by 45 cycles of 95°C for 10 s and 60 °C for 1 min. β-actin was used as the internal control. Each experiment using this analysis was replicated thrice.

### Immunoprecipitation

Cells were washed in PBS and lysed in lysis buffer A (25 mM Tris-HCl [pH 7.5], 420 mM NaCl, 0.5% Triton X-100, 5 mM CaCl_2_, 10 μg/mL aprotinin, 10 μg/mL leupeptin, and 1 mM phenylmethylsulfonyl fluoride [PMSF]) for 30 min at 4°C. Lysates were diluted 1.8-fold with lysis buffer B (25 mM Tris-HCl [pH 7.5], 0.5% Triton X-100, 5 mM CaCl_2_, 10 μg/mL aprotinin, 10 μg/mL leupeptin, and 1 mM PMSF) and were then centrifuged at 15,000 rpm for 10 min. Immunoprecipitation assays were performed using nProtein A Sepharose 4 Fast Flow or nProtein G Sepharose 4 Fast Flow (GE Healthcare) according to the manufacturer’s instructions. Briefly, each sample was incubated for 1 h at 4°C with anti-MTA1 antibody (Abcam), anti-S100A4 antibody (Developmental Studies Hybridoma Bank, Iowa City, IA, USA), normal rabbit IgG (Cell Signaling Technology), or normal mouse IgG (Sigma-Aldrich). Immune complexes were recovered on nProtein A Sepharose beads or nProtein G Sepharose beads for 1 h at 4°C. The immunoprecipitates were washed five times with TBS (50 mM Tris-HCl [pH 7.5], 150 mM NaCl), and 2× additional buffer (1 M Tris-HCl [pH 6.8], 10%SDS, 100% glycerol, bromophenol blue) was added. After boiling for 3 min at 95°C, the supernatants were used as western blotting samples. The experiment was repeated four times.

### Statistical analyses

The number of samples to be analyzed was predetermined based on the requirement of the statistical test used. Statistical analyses were conducted using Student’s *t*-tests to evaluate *MTA1* mRNA expression and NMIIa phosphorylation. Student’s *t*-tests or Welch’s *t*-tests were applied for the evaluation of tube formation assays (after the equality of variance was judged using the F-test) and the differences between two or multiple groups. Differences with P values < 0.05 were considered significant. The data points in the present study were not excluded. The researchers involved were not blinded, and the samples were not randomized during sample collection and data analysis.

## Data Availability Section

This study includes no data deposited in external repositories.

## Acknowledgements

We are grateful to Dr. K. Takenaga (Chiba Cancer Center Research Institute, Chiba, Japan) for kindly providing the MTA1 expression vector.

## Author contributions

MI, MO, NU, TO, HK, and FO designed the experiments. MI, NU, TO, and MO performed the experiments and analyzed the data. MI, MO, NU, TO, HK, and FO wrote, reviewed, and revised the manuscript.

## Expanded View Figure Legends

**Fig EV1** Generation of MTA1-KO MSS31 cell lines using the CRISPR/Cas9 system. (A) Western blotting of MTA1 protein levels in two MTA1-KO clones compared with those in wild-type MSS31 cells (n=3). (B) Alignment of the genomic region including the target sequence of gRNA in the genomic DNA of wild-type MSS31 cells, MTA1-KO clone #1, and MTA1-KO clone #2. The arrow represents the gRNA binding site. The first ATG codon is surrounded by a rectangle in the sequence gRNA: guide RNA

**Fig EV2** MTA1 was overexpressed in MTA1-KO MSS31 cells transfected with the MTA1 expression vector. (A) Expression of *MTA1* mRNA in wild-type MSS31 cells, MTA1-KO MSS31 cells (Non-treatment), MTA1-KO cells transfected with empty vector (EV), and MTA1-KO cells transfected with MTA1 expression vector (MTA1-KO+OE MSS31 cells: OE) analyzed by real-time PCR (n = 3). (B) Typical expression of MTA1 protein in wild-type MSS31 cells, MTA1-KO MSS31 cells (Non-treatment), MTA1-KO cells transfected with empty vector (EV), and MTA1-KO cells transfected with MTA1 expression vector (MTA1-KO+OE MSS31 cells: OE analyzed by western blotting (n = 3)

**Fig EV3** VEGFR2 was phosphorylated by VEGF and its expression was not altered by *MTA1* knockout. Phosphorylated-VEGFR2 (p-VEGFR2), VEGFR2 expression in wild-type MSS31 cells and MTA1-KO clones treated with VEGF (10 ng/mL) for 30 min was assessed by western blotting. Phosphorylated VEGFR2 and total VEGFR2 expression levels were determined by western blotting with the respective antibodies

**Fig EV4** MTA1 expression was not altered by VEGF stimulation. Western blotting for MTA1 expression in cells treated with VEGF (0, 10, 50 ng/mL) for 24, 48, and 72 h

